# Exacerbation of neurodegeneration by a metabolically engineered bacterium in a mouse model of tauopathy

**DOI:** 10.64898/2026.09.28.755025

**Authors:** Wade K. Self, Prabal Sharma, Grace Gent, Megan E. Bosch, Yiyang Zhu, Noah Powell, Cansu Agca, Jason D. Ulrich, Jianxun Song, Paul de Figueiredo, David M. Holtzman

## Abstract

**Background:** Therapeutics that modulate peripheral and central immune responses are emerging as promising approaches for neurodegenerative disease. However, the biological context in which these interventions are deployed may critically influence therapeutic outcomes. An attenuated *Brucella melitensis* 16M strain and a similar strain engineered to express the metabolite indole have previously demonstrated therapeutic efficacy in preclinical models of cancer and autoimmunity through immune system modulation. Here, we tested whether repeated administration of these bacterial strains could modify pathological outcomes in a mouse model of tauopathy.

**Methods:** Attenuated *Brucella* strains were administered to transgenic mice overexpressing mutant tau protein on a human APOE4 knock-in background via intraperitoneal injection every 5 weeks starting at 6 months of age. Mice were analyzed at 9.5 months of age for tau pathology, neurodegeneration, and changes in innate and adaptive immune responses.

**Results:** *Brucella* strains distributed predominantly in the periphery following repeated intraperitoneal treatment in transgenic mice, with minimal detection in the brain and no detection in the dural meninges. Repeated administration of either bacterial strain was well tolerated, with no evidence of adverse safety events. Surprisingly, repeated treatment with the engineered *Brucella* strain exacerbated neurodegenerative pathology, resulting in reduced hippocampal volume and increased phosphorylated tau pathology compared with control- and non-engineered *Brucella* strain -treated animals. Immunohistochemical analyses revealed increased numbers of hippocampal CD4^+^ T cells following treatment with either *Brucella* strain relative to PBS controls. However, only treatment with the engineered *Brucella* increased microglial reactivity within the hippocampus.

**Conclusion:** These findings demonstrate that peripheral administration of an engineered bacterial therapeutic can differentially influence neuroimmune responses and disease progression. These results underscore the importance of disease context when developing bacteria-based immunomodulatory therapies for neurodegenerative disorders.

## Background

Evidence from human genetic studies(1) and mouse models(2,3) suggest that microglia-mediated neuroinflammation promotes neurodegeneration caused by accumulation of misfolded microtubule-associated protein tau (tau) that is a posited cause for a class of neurodegenerative diseases known as tauopathies, which includes Alzheimer’s Disease (AD). The recent discoveries of T cells in the brain parenchyma of individuals with AD and primary tauopathies post-mortem (4–6) and protection from tau-mediated neurodegeneration after T cell or conventional type 1 dendritic cell depletion in mice (5,6) implicate adaptive immunity, and cells that reside outside of the central nervous system, in this process. Combined with observations that accumulation of tau pathology more directly correlates with regional brain atrophy and cognitive/functional decline in AD than amyloid plaque pathology(7,8), these observations suggest that targeting regulators of innate and adaptive immunity are a viable therapeutic strategy for neuroprotection in tauopathy, including AD. However, while immunotherapies currently exist for other disorders, there is an incomplete understanding of the appropriate mechanisms to target immune cell function in tauopathy.

A promising strategy for immunotherapies is the administration of attenuated, antigen-expressing pathogens such as viruses or bacteria to stimulate the immune system for disease modification. Such an approach has garnered significant attention recently based on epidemiological and “natural” experiment studies suggesting vaccines against herpes zoster virus and varicella zoster virus are associated with a lower risk of dementia in multiple human cohorts(9–13). A prominent example of an established treatment with live, attenuated bacteria is the Bacille Calmette-Guiren vaccine, a strain of *Mycobacterium bovis* originally developed for Tuberculosis vaccination that is also an established treatment for bladder cancer(14) and being investigated for the treatment of autoimmune diseases like Type 1 diabetes (NCT02081326, NCT0610891). With similar rationale, recent studies with an attenuated strain of *Brucella melitensis* 16M harboring a deletion in vjbR, a master regulator of virulence (BmΔvjbR), demonstrated anti-tumor efficacy in a murine colon adenocarcinoma model(15) and further engineering of a BmΔvjbR strain to produce indole via expression of tryptophanase A (BmΔvjbR::tnaA) reduced autoimmune responses in a murine collagen-induced arthritis model(16). Despite these promising results, whether such strategies can be deployed for the treatment of progressive neurodegenerative conditions like tauopathy has been relatively underexplored.

In this investigation, we tested whether repeated administration of attenuated *Brucella* strains BmΔvjbR and BmΔvjbR::tnaA would therapeutically modify neurodegenerative phenotypes in a murine model of tauopathy. Our initial hypothesis was that BmΔvjbR::tnaA would demonstrate therapeutic benefit via promotion of anti-inflammatory immune responses in the brain similar to phenotypes observed in the collagen-induced arthritis model (16). To our surprise, we observed an exacerbation of tau-mediated neurodegeneration with BmΔvjbR::tnaA treatment, highlighting the importance of disease context for the development of bacteria-based immunotherapies for neurodegeneration.

## Methods

### Animals and Study Design

All animal procedures and protocols were approved by the Institutional Animal Care and Use Committee at Washington University School of Medicine (IACUC: protocol number 24-0098). Male PS19 tau transgenic mice overexpressing human P301S tau by a mouse prion protein promoter (17) on a human APOE4 knock-in background(18) were generated by breeding males expressing the mutant tau transgene (TE4) to non-transgenic female mice (E4). All mice were housed in specific pathogen-free conditions, maintained under a consistent 12-h light/12-h dark cycle, with ambient room temperatures ranging from 20 °C to 26 °C and humidity levels between 30% and 70%. Food and water were available ad libitum.

Two studies were primarily performed in male TE4 mice: an assessment of *Bm*Δ*vjbR* bacterial distribution in tissues upon intraperitoneal injection and determining how repeated treatment of *Bm*Δ*vjbR* strains starting at 180 days of age would impact neuropathological outcomes at 285 days of age. The timepoints for treatment and assessment and the number of animals used for each treatment group were decided based on previous experience with these bacterial strains and tauopathy model by the investigators(5,6,15,16,19). Animals were randomly assigned to treatment groups. A small group (n = 2-5) of non-transgenic, male E4 mice were also treated and serve as qualitative references for various measurements, but studies were not powered to include these animals in statistical analyses. All animals that reached the 285-day end point were assessed for hippocampal and entorhinal/piriform cortex volume, ventricular volume, and immunohistochemical measures. For assessments of plasma metabolites and cytokines, samples selected for analysis were chosen based upon proximity to the mean measure of hippocampal volume within each treatment group. For each assay employed, scientists that processed the tissue and analysts that acquired and quantified images were blinded to the experimental conditions for a sample (i.e. animal genotype, treatment group).

### Brucella Melitensis 16M ΔvjbR (BmΔvjbR) Bacterial Culture and Treatment

Cultures were prepared similar to methods described previously (15). Briefly, *Bm*Δ*vjbR* strains were cultured from a 30% glycerol stock on tryptic soy agar (TSA) plates and incubated at 37°C for 3–4 days. *Bm*Δ*vjbR* individual colonies were inoculated into tryptic soy broth (TSB) and grown at 37°C with shaking (180–220 rpm) for 24–48 hour to generate a 5 mL preculture. For expansion, precultures were used to inoculate fresh TSB at 1–5% (v/v) of the final culture volume and incubated at 37°C with shaking (180–220 rpm) for an additional 24–48 hours. The same procedure was performed for *Bm*Δ*vjbR::tnaA* strain with TSA plates and TSB supplemented with chloramphenicol at 30 ug/mL. Bacterial growth was monitored spectrophotometrically by measuring optical density at 600 nm (OD600). An OD600 of 0.4 corresponded to approximately 1 × 10^9^ colony-forming units (CFU)/mL (Additional Figure 1A-B). Exponentially growing cultures (OD600 = 0.4–1.0) were used for *in vivo* administration.

For animal inoculations, bacterial cultures were harvested by centrifugation at 5,000 × g for 5 min at 4°C. Pelleted bacteria were washed three times with sterile phosphate-buffered saline (PBS; pH 7.2), with centrifugation at 5,000 × g for 5 min at 4°C between washes. Following the final wash, bacterial pellets were resuspended in sterile PBS and maintained on ice until administration. Bacterial concentrations were adjusted based on OD600 measurements to achieve the desired dose of 1.0e8 CFU/animal. The final concentration of each inoculate was confirmed via plating of diluted samples on TSA plates followed by CFU counting (Additional Figure 1C).

Animals were treated at designated time points via 100 uL intraperitoneal injection of either PBS, *Bm*Δ*vjbR,* or *Bm*Δ*vjbR::tnaA* with an insulin syringe (BD). Animals were monitored daily for 7 days after each inoculation.

### Nestlet Building Behavior

96 hours before euthanasia, group-housed mice were switched to individual housing. Pre-weighed nestlets (4 squares, 2.2-2.5g) were introduced into each cage. The next morning after 18 hours, A 5-point scale was used to score the results: the scoring system is based on the percentage of nesting material remaining plus the shredding conditions: score = 1, nest shredding < 25%; 2, nest shredding 25%–50%; 3, nest shredding 50%–90%; 4, nest shredding > 90%, but nest was not compacted; 5, complete nest built(20).

### Tissue Harvesting

Prior to transcardiac perfusion, mice were anesthetized with pentobarbital (50 mg/kg, intraperitoneal). Blood was collected from the heart before transcardiac perfusion, which was centrifuged at 5000xg for 5mins at 4°C to obtain plasma. Mice were transcardially perfused with ice-cold phosphate-buffered saline containing 0.3% heparin.

For animals in which bacterial enumeration was performed, spleen, dural meninges, and whole brain were dissected and placed in cold PBS. Samples were weighed and homogenized in 1 mL PBS. A 1:25 dilution of spleen, a 1:2 dilution of meninges, and a 1:2 dilution of whole brain was plated on TSA plates and CFU counted 96 hours later.

For animals in the therapeutic cohort, one hemibrain was dissected, snap-frozen and stored at −80°C for biochemical analyses. The other hemibrain was immerse-fixed for 24h in 4% paraformaldehyde following by cryoprotection in 30% sucrose for 48h prior to sectioning Brains were cut coronally into 50 μm-thick sections on a freezing sliding microtome (Leica SM1020R) and stored in cryoprotectant solution (0.2 M PBS, 15% sucrose, 33% ethylene glycol) at −20 °C until use in immunohistochemical analyses.

### Volumetric Analysis

Volumetric analysis of the hippocampus, entorhinal/piriform cortex (EC/PC), and ventricle was performed via stereological methods by assessing sections spaced by 300μm starting from bregma −1.1 mm to bregma −4.1 mm as described previously (19,21). Briefly, 50μm microtome-cut sections mounted on slides were briefly immersed in distilled water before incubating in pre-warmed 0.1% cresyl violet acetate solution at for six minutes. Tissues were rinsed in distilled water briefly and transferred to 70%, 95% and 100% ethanol sequentially, for two minutes each. Slides were then cleared in xylene before coverslipping with Cytoseal60 mounting medium (Thermo Fisher Scientific). Slides were scanned using Hamamatsu’s Nanozoomer microscope at 20X magnification. Hippocampus, EC/PC, and ventricles were traced using NDP.view 2. The formula for the volumetric calculation was volume = (sum of area) * 0.3 mm.

Hippocampal volume was calculated for all animals analyzed. In some cases, animals were excluded from analysis for EC/PC and lateral ventricle volume due to significant damage to the EC/PC upon mounting that compromised the ability to reliably quantify these areas.

### Neuronal Thickness Layer Measurements

Dentate granule and CA1 cell layers were measured across three sections corresponding to -2.3 mm, -2.6 mm, and -2.9 bregma by drawing a scale line across the cell layer using NDP.view 2 and calculating the average value for each mouse.

### Immunohistochemistry

Free-floating sections corresponding to -1.7 mm and -2.0 mm bregma were briefly washed with TRIS buffered saline (TBS) before quenching endogenous peroxidase with 0.3% hydrogen peroxide in TBS for 10 minutes at room temperature. After a brief wash in TBS, sections were blocked using 3% milk-0.1% TritonX100 (TBS-T) for 30 minutes at room temperature, followed by washing with TBS for 5 minutes three times at room temperature. Primary antibody incubation in biotinylated AT8 (Phospho-Tau Ser202, Thr205; 1:500, MN1020B, Thermo Fisher Scientific) was performed overnight at 4°C.

The following day, sections were briefly washed three times for 5 minutes with TBS before incubation with ABC Elite Solution (Vectastain PK-6100) for 60 minutes. Sections were washed three times in TBS for 5 minutes at room temperature before being developed with 3, 3’-diaminobenzidine (DAB, Sigma, D5905) for 8 minutes. Sections were washed three times for 5 minutes at room temperature and mounted on slides.

The following day, dried slides were dehydrated using a series of increasingly concentrated ethanol (50%, 70%, 95%, 95%, 100%, 100% for 1 minute each) before finally immersing in xylene twice for 4 minutes. Slides were immediately coverslipped using Cytoseal mounting medium.

### Immunofluorescence

Free-floating sections corresponding to -2.0 and -2.3 mm bregma were washed three times with TBS for 5 minutes at room temperature followed by permeabilization with TBS-T for 30 minutes.

Sections for assessing microgliosis were processed similarly to previous methods (6). Briefly, sections were blocked in 10% donkey serum in TBS-T for 30 minutes at room temperature. Primary antibodies were diluted in blocking buffer and incubated at 4°C overnight with slow agitation unless stated otherwise. Primary antibodies were used as follows: Iba1 (1:500; NB100-1028, Novus Biologicals), MHC-II (1:1000, BioLegend, 107603), P2RY12 (1:1000, AnaSpec, AS-55043A). The following day, sections were washed and incubated with secondary antibody diluted in blocking buffer (Donkey-anti-goat IgG (H+L)-AlexaFluor647, 1:500, Thermo Fisher Scientific; Donkey-anti-rabbit IgG (H+L)-AlexaFluor568, 1:500, Thermo Fisher Scientific; Donkey-anti-rat IgG (H+L)-AlexaFluor488, 1:500) followed by incubation with 4′,6-Diamidin-2-phenylindol (DAPI 1:10,000) for 5 minutes before mounting sections onto slides (Prolong^™^ Gold Antifade reagent, Thermo Fisher Scientific).

Sections for assessing T cell abundance were similar to methods described previously (6). Briefly, sections were blocked with a solution containing 10% normal donkey serum, 1% bovine serum albumin (BSA), and RecombiMAb anti-mouse CD16/CD32 FcR blocker (BioxCell, CP025; 1:400) in TBS buffer with 0.5% Triton X-100 for 2 hour at room temperature, followed by three days incubation at 4°C with primary antibodies rat anti-CD3ε (Leinco Technologies, C2457 ; 1:400), human anti-CD4 (Miltenyi Biotec, 130-123-215; 1:200), and rabbit anti-CD8α (Invitrogen, MA5-29682; 1: 200). After three days incubation, brain sections were then washed in TBSx for three times (10 mins/cycle), followed by incubation with fluorescent-labeled secondary antibodies (Thermo Fisher Scientific, 1:500) for 2 hours at room temperature. Sections were washed with TBSx followed by incubation with DAPI for 5 minutes (1:10,000 at room temperature). Sections were counterstained with TrueBlack Lipofuscin Autofluorescence Quencher (Biotium 23007) for 1 minute, quickly washed in TBS and mounted in ProLong Diamond Antifade mounting medium (Invitrogen, P36961).

### Imaging and Quantification

For AT8 DAB imaging, slides were scanned using Hamamatsu’s Nanozoomer microscope at 20X magnification. For assessments of microgliosis, 20 um z-stack images (5 um/stack) of the entire hippocampus were acquired on a Leica THUNDER Imager Tissue at 20X magnification processed with the Instant Computational Clearing Algorithm. For assessments of T cell abundances, 20 um z-stack images (1.5 um/stack) of the entire hippocampus were acquired on a Leica Stellaris 5 confocal microscope at 20X magnification and 512^512 resolution.

AT8 DAB hippocampal images were exported from NDP.view 2 and converted to 8-bit images in ImageJ (NIH v1.53). For assessments of microgliosis, maximum projections of the individual channels for merged hippocampal images were exported for percent area analysis for each marker. For both AT8 and microgliosis, regions of interest (ROI) containing the hippocampus were traced for each section, and the percentage area of each marker within an ROI was calculated using a predetermined threshold for each stain with the Adjust Threshold function in ImageJ.

For quantification of T cell numbers, maximum projections of the merged channel scaled viewer image or each hippocampus was exported into ImageJ. Manual counting of cells weas performed in ImageJ by an analyst blindly labeled to experimental conditions. CD8^+^ T cells were counted based on colocalization of CD3, CD8, and DAPI signal. CD4^+^ T cells were counted based on colocalization of CD3, CD4 and DAPI signal.

All data points plotted represent the average of the two tissue sections analyzed per mouse. All representative images reflect a data point near the average value for each treatment group.

### Plasma Untargeted Metabolomics

50 uL of mouse plasma samples were subjected to metabolite extraction using 80% Methanol with three rounds of vortex mixing followed by overnight incubation. The supernatant containing metabolites was centrifuged, collected, and then dried using a SpeedVac with no heat. The dried pellet was reconstituted in 50 µL of 50% MeOH, and subsequently injected into the LC-MS/MS system. The samples were subject to two liquid chromatography conditions on a Vanquish Horizon UHPLC system (Thermo Scientific): a C18 column or an Amide column. Mass Spectrometry was performed on an an Orbitrap Tribrid ID-X in both Positive and Negative ion mode with AcquireX Deep Scan (Thermo Scientific). Metabolite abundances were quantified by area under the curve based on MS1 intensity peaks, and log-transformed values underwent quantile normalization to limit technical variation and batch effects across analytical runs.

Blinded group identification numbers were used for initial statistical analysis. Pathway and chemical class enrichment tests were performed between each group based on the significant features in each comparison. Data points plotted in Figures represent the column and ionization mode where the highest area under the curve was observed for each metabolite of interest.

### Plasma Cytokine Immunoassay

25 uL of mouse plasma were analyzed with the ProcartaPlex Mouse Immune Monitoring Panel 48plex (Thermo Fisher Scientific, EPX480-20834-901) to measure cytokines and chemokines according to the manufacturer’s instructions(22). Belysa Software v1.2.1 (Millipore Sigma) was used to calculate absolute concentrations using a 5-parameter curve fit algorithm. Samples were run in duplicate on immunoassay plate and excluded from analysis if measurements were outside of the upper or lower limit of quantification for the standard curve. Outliers were identified using ROUT analysis with a Q value of 0.1% and excluded from analysis.

### Statistical Analysis

All statistical analyses were performed in GraphPad Prism 11. Categorical variables were assessed with Chi-Square contingency test. All remaining statistical tests involved analysis of three or more groups using analysis of variance testing for parametric and non-parametric data. The Shapiro-Wilk test was used for assessments of normality, and the Brown-Forsythe method was used to determine homoscedasticity of variance for a given dataset. Final statistical tests employed for each dataset are described in the Figure legends.

## Results

### BmΔvjbR strain distribution and safety profile in TE4 mice

*Brucella* infections in humans can manifest in neurological symptoms termed neurobrucellosis(23). Therefore, we first sought to understand if the *Bm*Δ*vjbR* strain could be found in the brain if administered peripherally and assess the tolerability of *Bm*Δ*vjbR* strain treatment in our therapeutic cohort. To test this, we first performed repeated intraperitoneal injections of either vehicle (PBS) or *Bm*Δ*vjbR::tnaA* in 150-day-old TE4 mice followed by harvesting tissues 24 hours after the final treatment at 250 days (Figure 1A). *Bm*Δ*vjbR::tnaA* colony-forming units (CFU) were observed in splenic tissue (357 ± 320 CFU/mg), with little to no detection in whole brain or dural meningeal lysates (Figure 1B). In our therapeutic cohort (Figure 2), incidence of premature deaths or end-stage body weight in TE4-treated animals did not differ by experimental group (Figure 2B,C).

**Figure 1.**
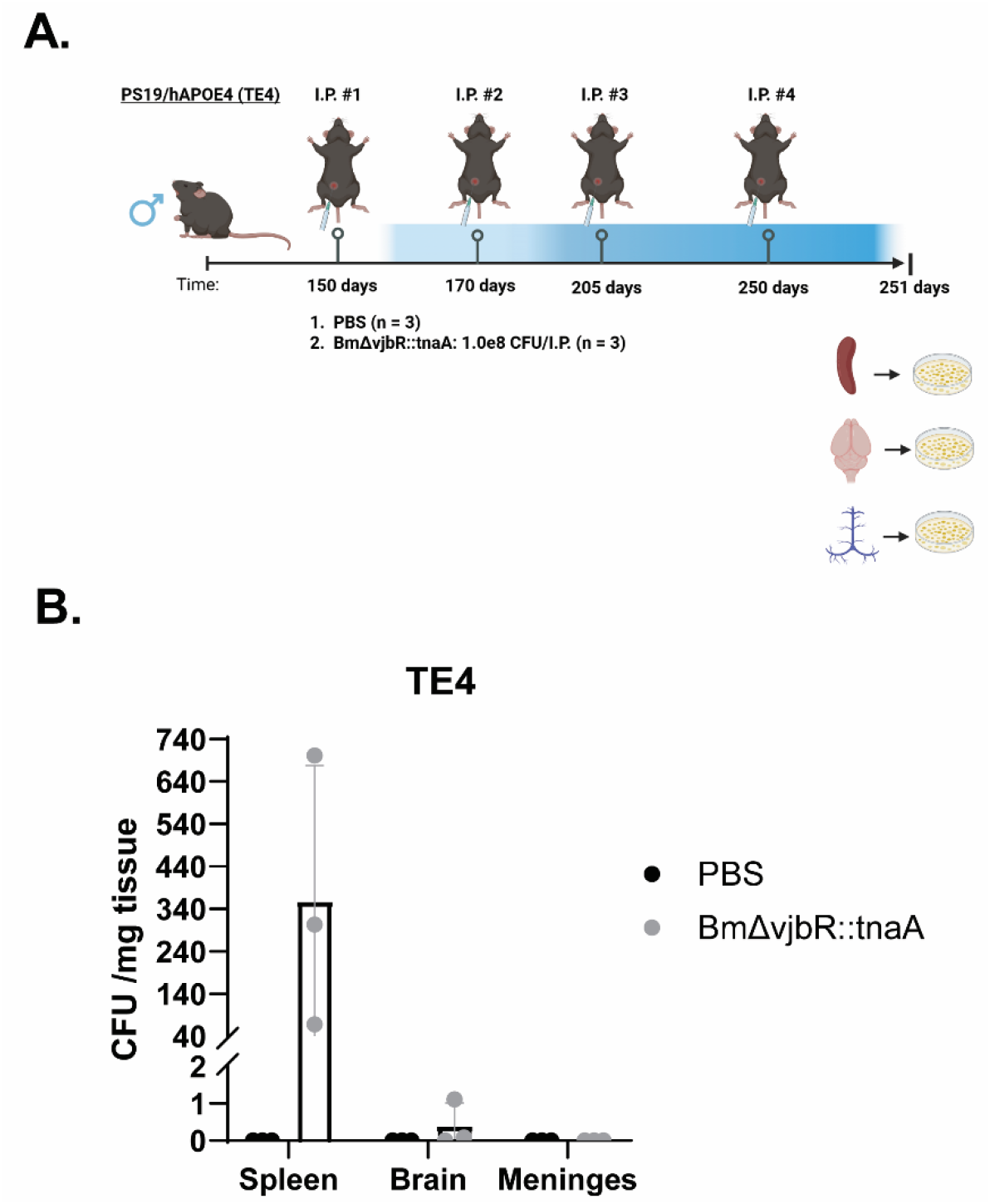
BmΔvjbR::tnaA distributes primarily in the periphery after repeated treatment. A. Experimental design to determine abundance of BmΔvjbR::tnaA after repeated dosing in TE4 male mice. B. Quantification of Colony Forming Unit (CFU) assay on various tissues. Individual data points correspond to one mouse. Bar height represents mean value; error bars represent standard deviation. Illustrations in A. created with Biorender.

**Figure 2.**
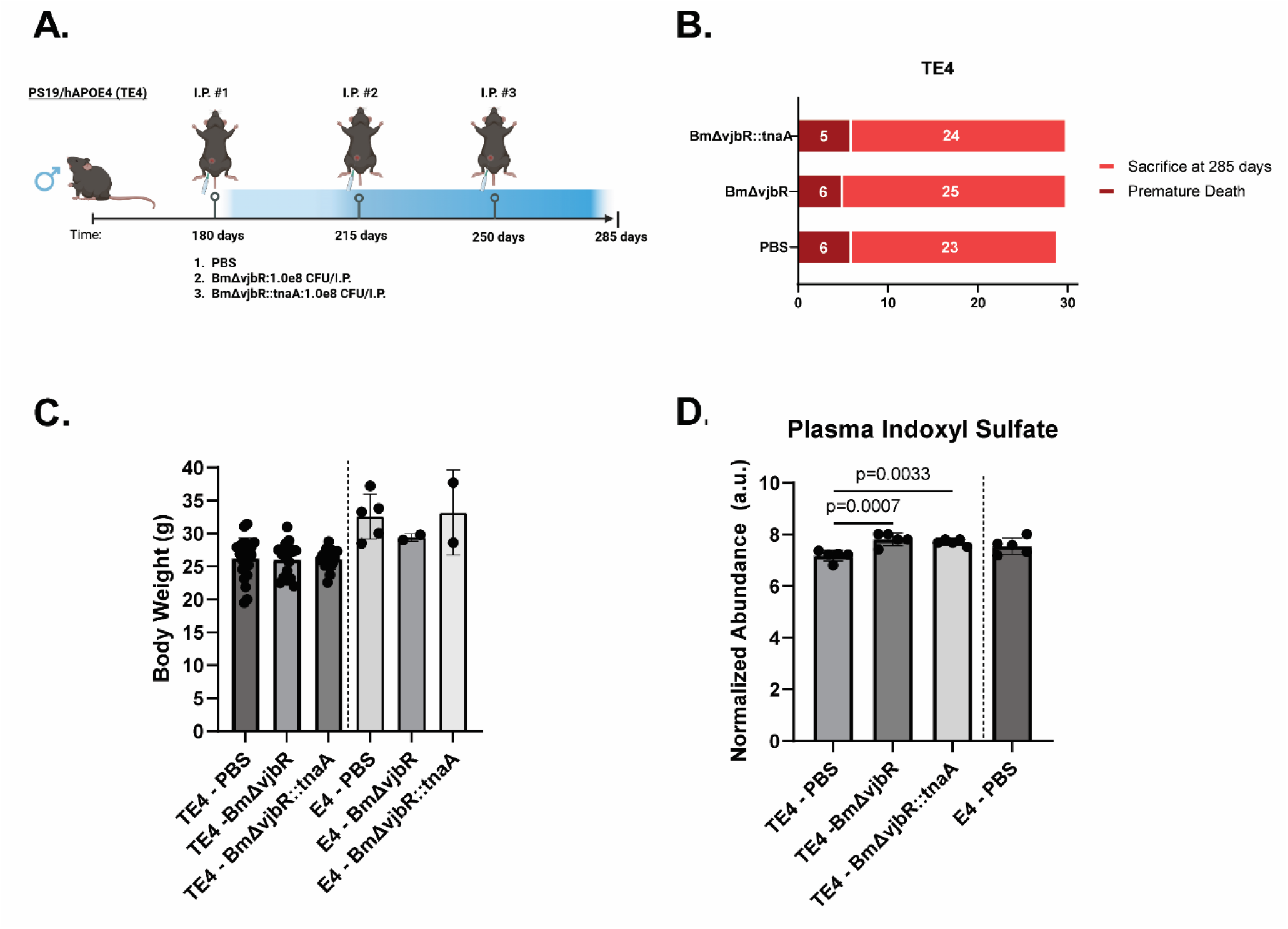
Experimental design and assessments of safety in TE4 therapeutic cohort. A.) Experimental design to assess effects of *Brucella* strains on tau-mediated neurodegeneration in TE4 male mice. Animals received first treatment at 180 days of age and received the same treatment every 5 weeks via intraperitoneal injection (I.P.) until sacrifice at 285 days of age. B.) Survival outcomes of TE4-treated animals. Χ^2^: 0.1781, p-value: 0.91. C.) End-stage body weight of mice treated in therapeutic cohort. 1-way ANOVA on TE4-treated animals: F: 0.02162 (2,53), p-value: 0.98. D.) Quantile-normalized abundances of plasma indoxyl sulfate levels in end-stage, 285-day old mice. 1-way ANOVA on TE4-treated animals: F: 14.71 (2,12), p: 0.006. N = 5 animals per TE4-treated group for metabolomic analysis. P-values for individual comparisons with Tukey’s multiple comparisons test *post hoc*. Bar height represents mean value; error bars represent standard deviation. Illustrations in A. created with Biorender.

In normal physiological conditions, indole is predominantly produced via catabolism of dietary tryptophan by microbiota and is further transformed in the liver into indoxyl sulfate, a molecule that is a known uremic toxin when clearance is impaired (24). We performed plasma metabolomics analysis to test whether indoxyl sulfate or other tryptophan catabolites were altered due to *Bm*Δ*vjbR::tnaA* treatment in our therapeutic cohort. Elevated levels of plasma indoxyl sulfate were observed between PBS-treated and *Bm*Δ*vjbR* strains, but no differences in relative abundance were observed between *Bm*Δ*vjbR::tnaA*-treated and *Bm*Δ*vjbR*-treated TE4 mice (Figure 2D) or other measures of microbial-derived catabolites of tryptophan (Additional Figure 2). These findings were also reflected in pathway enrichment analysis, as there was no significant association between *Bm*Δ*vjbR::tnaA*-treated and *Bm*Δ*vjbR*-treated TE4 mice with pathway terms “Absorption, and excretion pathway of gut-derived uremic toxins” and “Metabolic pathway of tryptophan” (Additional Data File 1).

In sum, these results suggested that our dosing regimen resulted in distribution of *Bm*Δ*vjbR* strains outside of the central nervous system, and that *Bm*Δ*vjbR* administrations were well-tolerated in TE4 mice.

### Repeated BmΔvjbR::tnaA exacerbates tau-mediated neurodegeneration in TE4 mice

285-day-old male TE4 mice in the PBS-treated group experienced marked tissue atrophy and accumulation of hyperphosphorylated tau aggregates in the hippocampus and entorhinal/piriform cortex and dilation of the lateral ventricle compared to E4 mice that did not harbor the mutant tau transgene (Figure 3A), similar to previous reports(19,25). Significant increases in hippocampal and entorhinal/piriform cortical tissue atrophy were associated with *Bm*Δ*vjbR::tnaA* treatment compared to *Bm*Δ*vjbR-* treated TE4 mice with a trend for similar increases compared to PBS-treated TE4 mice (Figure 3B-D). Similarly, *Bm*Δ*vjbR::tnaA-*treated TE4 mice exhibited a trend toward lower neuronal cellular layer thicknesses in the Dentate Gyrus and Layer CA1 of the hippocampus compared to *Bm*Δ*vjbR-*treated TE4 mice (Additional Figure 3A-C). TE4 mice exhibit deficits in nestlet building behavior that correlates with brain atrophy, and *Bm*Δ*vjbR::tnaA*-treated TE4 mice had a lower median nestlet-building score compared to *Bm*Δ*vjbR-* or PBS-treated animals, although these results did not reach statistical significance (Additional Figure 3D).

**Figure 3.**
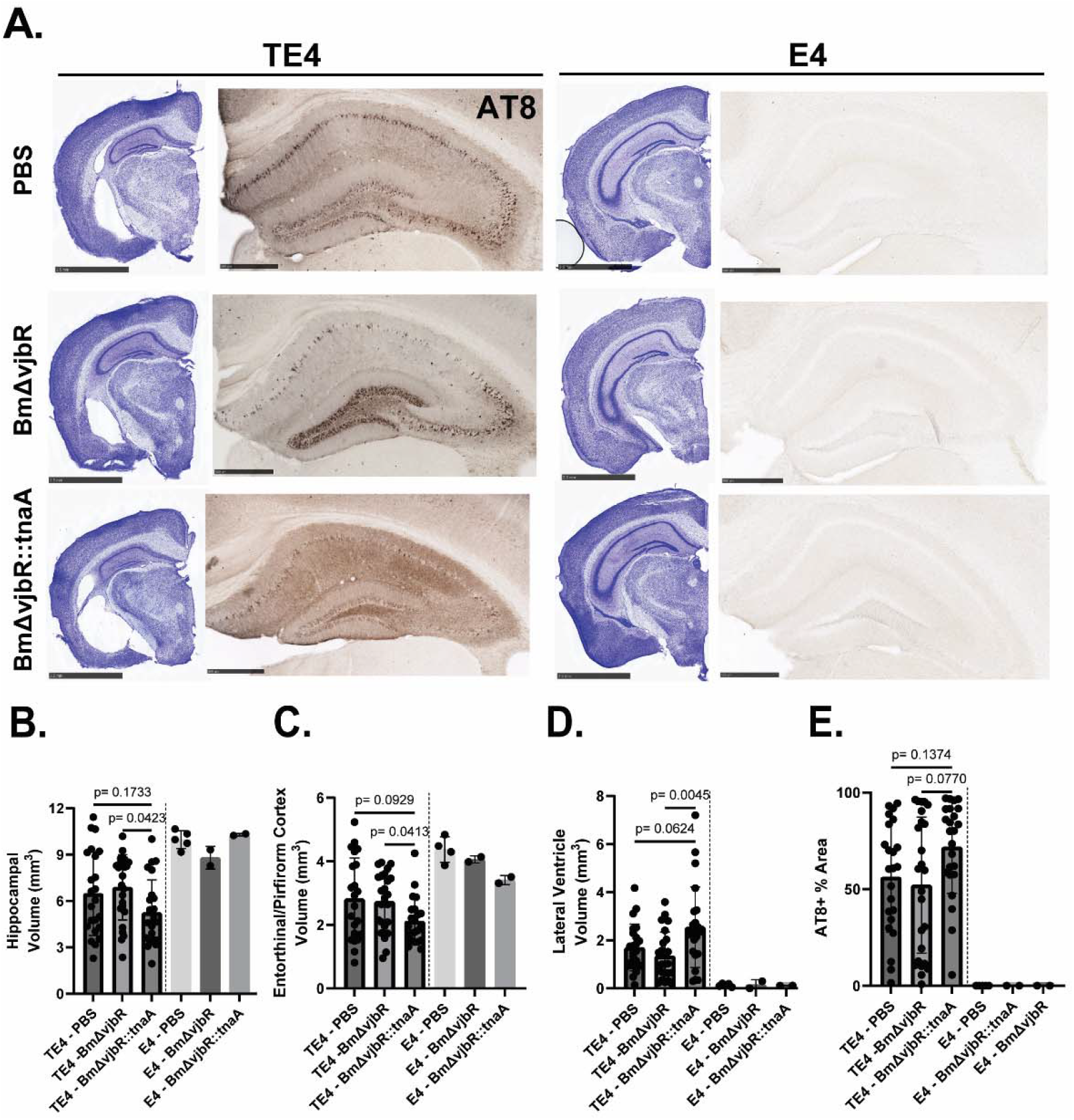
Repeated BmΔvjbR::tnaA treatment exacerbates tau-mediated neurodegeneration. A.) Representative images of tissue atrophy and AT8+ tau accumulation in the hippocampus of treated TE4 and E4 animals at 285 days. Scale bar for Cresyl Violet: 2.5 mm. Scale bar for AT8: 500 um. B.) Volumetric analysis of hippocampus. F: 3.261 (2,69), p-value: 0.0443. Number of animals analyzed in TE4 groups: PBS: 23, BmΔvjbR: 25, BmΔvjbR::tnaA: 24. C.) Volumetric analysis of entorhinal/piriform cortex. W: 4.336 (2.000, 41.39), p-value: 0.0195. Number of animals analyzed in TE4 groups: PBS: 22, BmΔvjbR: 24, BmΔvjbR::tnaA: 22. D.) Volumetric analysis of lateral ventricle volume. F: 5.712 (2,68), p-value: 0.0051. Number of animals analyzed in TE4 groups: PBS: 23, BmΔvjbR: 24, BmΔvjbR::tnaA: 24. E.) Percentage AT8-positive coverage analysis in the hippocampus. W: 3.409 (2.000, 45.15), p-value: 0.0418. Number of animals analyzed in TE4 groups: PBS: 23, BmΔvjbR: 25, BmΔvjbR::tnaA: 24. F statistic denotes 1-Way ANOVA test and W statistic denotes Welch’s ANOVA test for TE4-treated groups. P-values for individual comparisons on graphs represent results of Tukey’s multiple comparisons test or Dunnett’s T3 multiple comparisons test *post hoc*. Bar height represents mean value; error bars represent standard deviation.

*Bm*Δ*vjbR::tnaA*-treated TE4 mice also showed increased mean AT8+ area coverage in the hippocampus compared to *Bm*Δ*vjbR-* or PBS-treated animals that approached statistical significance (Figure 3E). These results were the opposite of our initial hypothesis, in that *Bm*Δ*vjbR::tnaA* treatment appears to promote damage in the setting of tau-mediated neurodegeneration.

### Differential CNS and peripheral immune responses to BmΔvjbR strains in TE4 mice

Microglial activation is required for tau-mediated neurodegeneration and can be exacerbated by infiltrating T cells into the brain parenchyma of TE4 mice (3,5), so we assessed markers microglial activation states in the hippocampus using IBA1, P2RY12 (homeostatic microglia), and MHC-II (activated, IFNγ-responsive microglia) (Figure 4). Consistent with assessments of neurodegeneration, we observed a significant increase in MHC-II expression in the hippocampus of *Bm*Δ*vjbR::tnaA*-treated TE4 mice compared to PBS-treated mice and a trend when compared to the *Bm*Δ*vjbR* group (Figure 4B). No differences in IBA1 or P2RY12 hippocampal area coverage were observed between TE4-treated groups (Figure 4 B,C), possibly due to a ceiling and floor effect of increased IBA1 expression and reduced P2RY12 expression, respectively, in TE4 mice compared to E4 mice (see also Additional Figure 4). Increases in MHC-II expression in the hippocampus of *Bm*Δ*vjbR::tnaA*-treated mice suggested that there may be differences in T cell abundance between TE4-treated groups.

**Figure 4.**
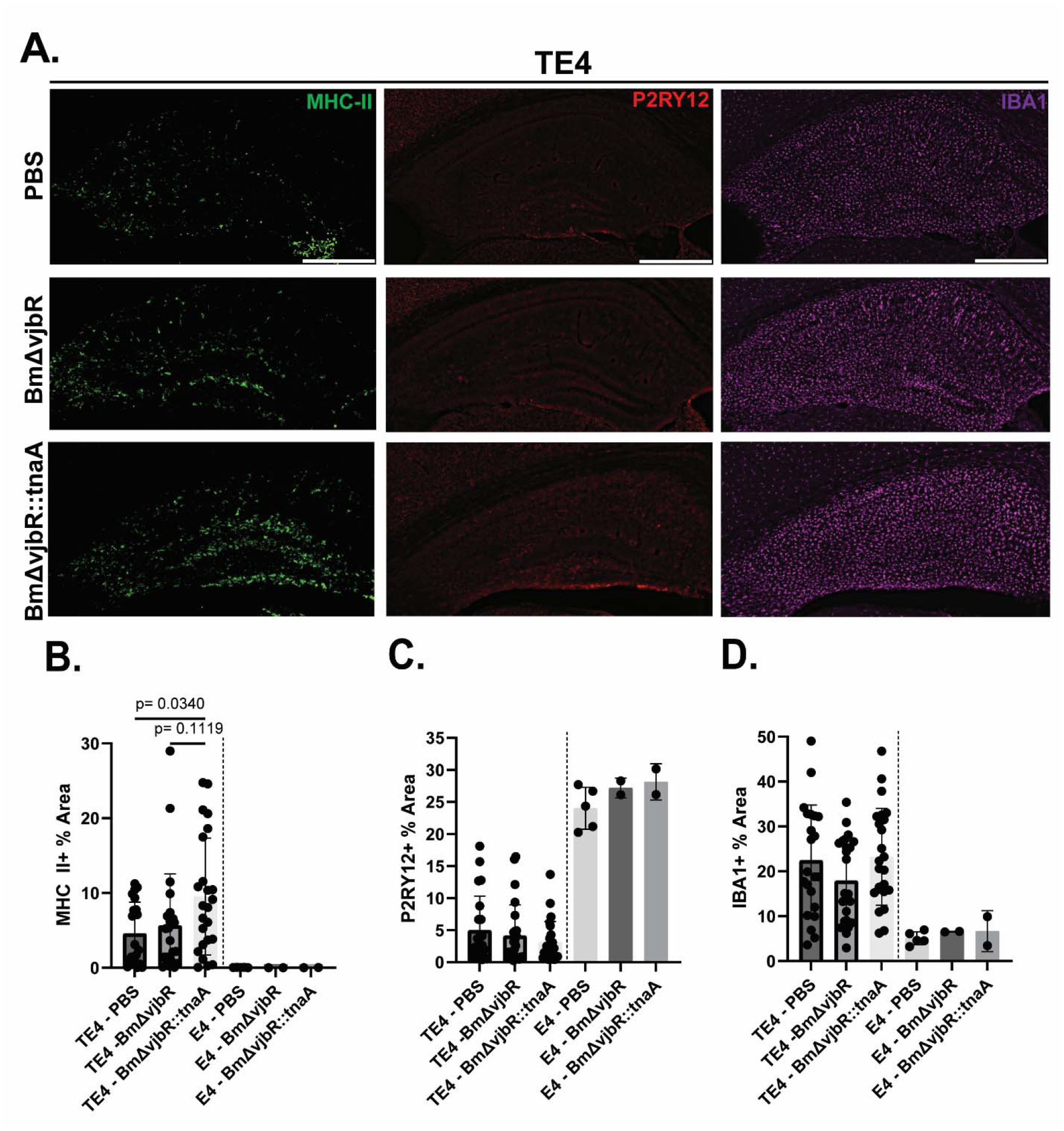
Repeated BmΔvjbR::tnaA treatment increases degeneration-associated microglial reactivity. A.) Representative images of hippocampal reactive microgliosis markers in 285-day-old TE4 mice. Scale bar: 500 um. B.) Percentage MHC-II-positive coverage analysis. F: 3.670 (2, 67), p-value: 0.0307. C.) Percentage P2RY12-positive coverage analysis. F: 1.102 (2, 67), p-value: 0.3690. D.) Percentage IBA1-positive coverage analysis. F: 1.655 (2,67), p-value: 0.1987. F statistics denote 1-Way ANOVA test on TE4-treated groups. P-values for individual comparisons on graphs represent results of Tukey’s multiple comparisons test *post hoc*. Bar height represents mean value; error bars represent standard deviation. Number of animals analyzed in TE4 groups: PBS: 22, BmΔvjbR: 24, BmΔvjbR::tnaA: 24.

Immunohistochemical analysis of T cell numbers in the hippocampus demonstrated a significant increase of CD4^+^ T cells in TE4 mice treated with either *Bm*Δ*vjbR* strain compared to PBS-treated animals (Figure 5B). A similar, smaller effect was observed in mean CD8^+^ T cells numbers that did not reach statistical significance (Figure 5C).

**Figure 5.**
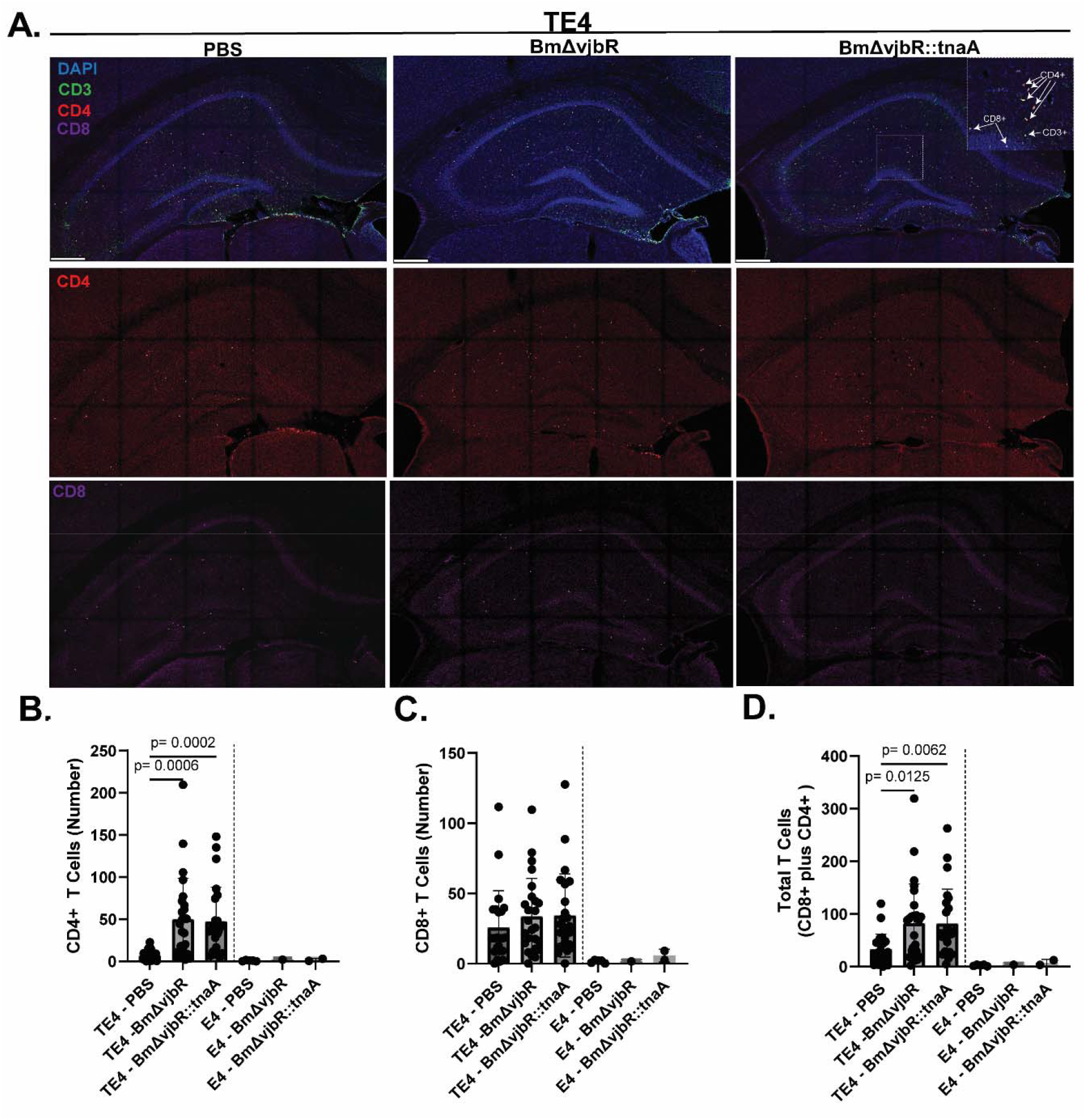
Repeated treatment with BmΔvjbR strains leads to increased CD4^+^ T cells in the hippocampus. A.) Representative images of T cell quantification in 285-day-old TE4 mice. Top is merged image of DAPI (blue), CD3 (green), CD4 (red), and CD8 (purple), and separate CD4 and CD8 channels. Enlarged section of BmΔvjbR::tnaA representative images provides example of colocalization of CD8 and CD4 with CD3. Scale bar: 250 um. B.) Quantification of hippocampal CD3^+^,CD4^+^ T cells. W: 20.39 (2, 32.48), p-value: <0.0001. C.) Quantification of hippocampal CD3^+^,CD8^+^ T cells. F: 0.7062 (2, 69), p-value: 0.4971. D.) Quantification of combined hippocampal CD4^+^ and CD8^+^ T cells. W: 8.832 (2.00, 39.80), p-value = 0.0007. F statistic denotes 1-Way ANOVA test and W statistic denotes Welch’s ANOVA test for TE4-treated groups. P-values for individual comparisons on graphs represent results of Tukey’s multiple comparisons test or Dunnett’s T3 multiple comparisons test *post hoc*. Bar height represents mean value; error bars represent standard deviation. Number of animals analyzed in TE4 groups: PBS: 23, BmΔvjbR: 25, BmΔvjbR::tnaA: 24.

Intriguingly, these data suggest that a primary consequence of repeated *Bm*Δ*vjbR* exposure in the periphery in the setting of tauopathy results in a preferential increase in CD4^+^ T cells in areas prone to degeneration.

These data prompted us to ask whether peripheral immune activation could explain the differential phenotypes observed in the brain, particularly between *Bm*Δ*vjbR and Bm*Δ*vjbR::tnaA* treatment. Spleen weights were significantly larger in both *Bm*Δ*vjbR* strain groups compared to TE4-treated animals, suggesting that the repeated dosing paradigm had lasting effects on the peripheral immune status of TE4-treated mice despite the last dose being administered 5 weeks before euthanasia (Figure 6A). We assessed the abundance of 48 plasma cytokines and observed significant changes in 3 molecules: IL-1β, IL-10, and IL-3, in TE4-treated animals (Figures 6B-D, Additional Figure 5). Plasma IL-3 levels were significantly decreased in *Bm*Δ*vjbR::tnaA-*treated TE4 mice and trending towards a decrease in *Bm*Δ*vjbR-*treated TE4 animals compared to the PBS-treated TE4 group. Similar trends were observed in plasma CCL3 and CCL5 concentrations, but these results did not reach statistical significance (Figure 6 I, J).

**Figure 6.**
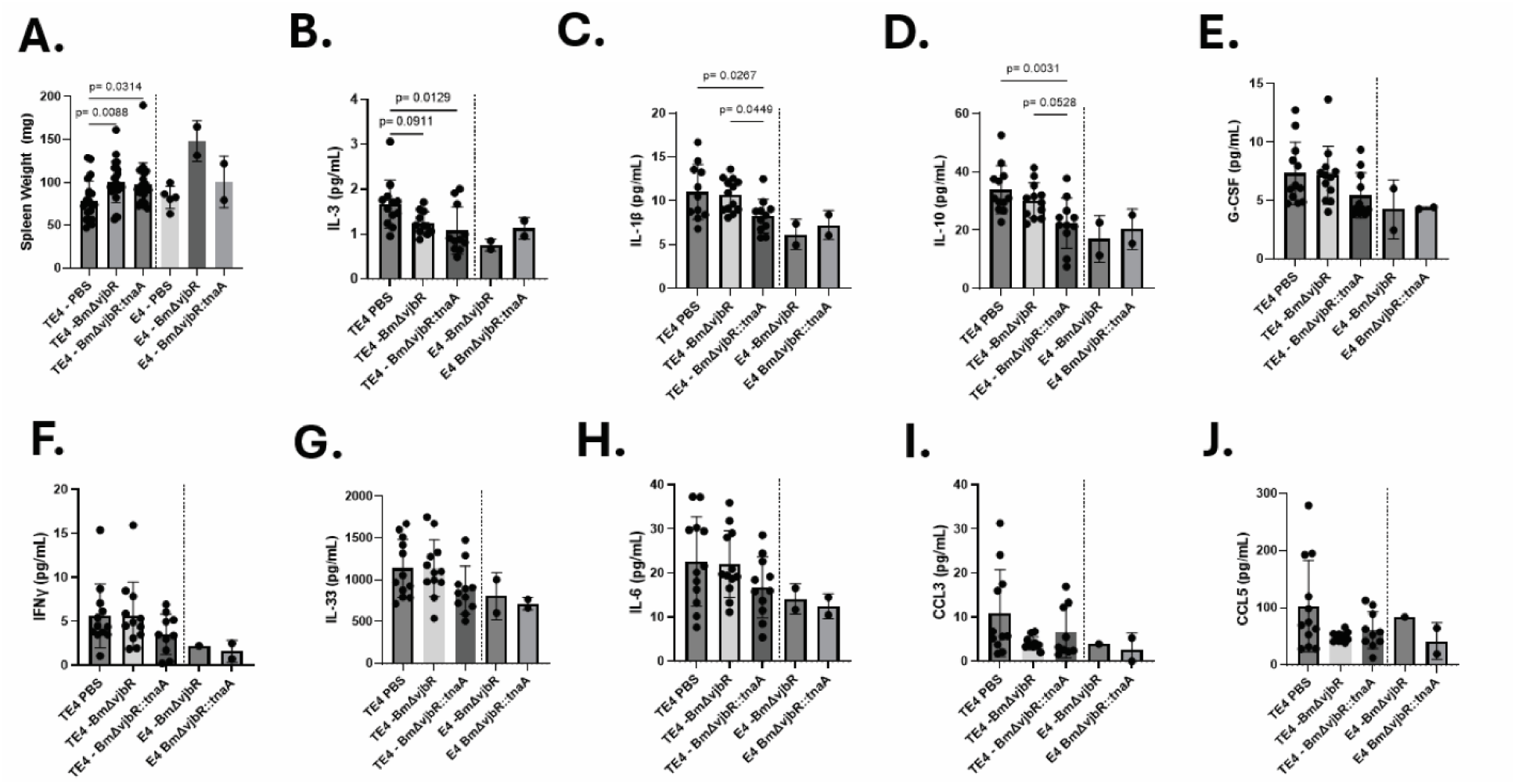
Repeated treatment with BmΔvjbR strains alters peripheral immune signatures. A.) Quantification of dissected spleen weights in 285-day-old animals at time of sacrifice. F: 5.395 (2, 63), p-value = 0.0069. B-H.) Quantification of select plasma cytokines and chemokines from the 48-plex immune monitoring panel, n = 10-12 per TE4-treated group. B.) IL-3 concentration. F: 4.957 (2,31), p-value: 0.0136. C.) IL-1β concentration. F: 4.538 (2,32), p-value: 0.0184. D.) IL-10 concentration. F: 6.647 (2,32), p-value: 0.0039. E.) G-CSF concentration: F: 2.136 (2, 32), p-value: 0.1347. F.) IFNγ concentration. F: 1.347 (2,31), p-value: 0.2749. G.) IL-33 concentration. F: 2.388 (2,32), p-value: 0.1080. H.) IL-6 concentration. F: 1.693 (2,32), p-value: 0.2001. I.) CCL3 concentration. W: 2.906 (2.000, 16.69), p-value: 0.0827. J.) CCL5 concentration. W: 2.975 (2.000, 14.15), p-value: 0.0834F statistic denotes 1-Way ANOVA test and W statistic denotes Welch’s ANOVA test for TE4-treated groups. P-values for individual comparisons on graphs represent results of Tukey’s multiple comparisons test or Dunnett’s T3 multiple comparisons test *post hoc*. Bar height represents mean value; error bars represent standard deviation.

Plasma IL-1β was significantly decreased in *Bm*Δ*vjbR::tnaA-*treated TE4 mice compared to *Bm*Δ*vjbR-*treated TE4 animals, and similar trends were observed for plasma IL-10. Similar observations of reduced plasma levels of G-CSF, IFNγ, IL-33, and IL-6 were observed in *Bm*Δ*vjbR::tnaA-*treated TE4 animals compared to the *Bm*Δ*vjbR-* treated TE4 group but did not reach statistical significance (Figure 6 E-H). Taken together, these data suggest that there may be differences in immune regulation in response to differential *Bm*Δ*vjbR* strain treatments which influence microglial-T cell interactions in the brain that contribute to tau-mediated neurodegeneration.

## Discussion

In this study, we found that repeated administration of a bacterial strain engineered to produce indole, BmΔvjbR::tnaA, results in divergent outcomes related to therapeutic modification in a model of tauopathy compared to previous results in a model of autoimmunity(16). Rather than reducing pathology as initially hypothesized, BmΔvjbR::tnaA treatment increased hippocampal and cortical atrophy, increased hippocampal MHC-II expression, and was associated with greater accumulation of hyperphosphorylated tau. These effects occurred without detectable bacterial colonization of the brain, in contrast to the mechanism of action of the parent BmΔvjbR strain in the context of anti-cancer immunity by homing to the tumor site to modify the inflammatory microenvironment (15). These observations were fortified by divergent results between animals treated with the BmΔvjbR and BmΔvjbR::tnaA, excluding the possibility that results are simply due to alterations in systemic inflammation due to bacterial infection. In line with this conclusion, plasma cytokine profiling in these animals demonstrated a lack of systemic hyperinflammation in mice treated with BmΔvjbR strains. Despite the manifestation of phenotypes opposite to therapeutic benefit in the BmΔvjbR::tnaA treatment group, these data support the idea that modulation of peripheral immune responses can influence neurodegenerative outcomes.

One of the most striking observations was the accumulation of CD4^+^ T cells in the hippocampus following repeated exposure to both BmΔvjbR strains. This finding suggests that chronic peripheral exposure to attenuated Brucella can reshape immune surveillance of the CNS during tauopathy such that certain T cell populations are more sensitive to signals promoting infiltration. Because increased CD4^+^ T-cell abundance was observed with both strains whereas worsened neurodegeneration was restricted primarily to the BmΔvjbR::tnaA group, T-cell recruitment alone is unlikely to explain the differential phenotype. Instead, our data suggests that the functional state of infiltrating T cells may be more important determinants of disease progression than absolute cell numbers. Consistent with this possibility, BmΔvjbR animals exhibited increased hippocampal MHC-II expression, but not overall levels of microgliosis as assessed by IBA1 area coverage within hippocampal tissue. MHC-II is induced on microglia during the late stages of neurodegeneration observed in the PS19/hAPOE4 mouse model, and possibly reflects interactions with infiltrating T cells, the primary source of interferon gamma production that induces MHC-II expression (5). Differences in MHC-II expression and neurodegeneration may point to differences in interferon gamma production between CD4 T cells treated with BmΔvjbR and BmΔvjbR::tnaA, raising the possibility of a divergence in the bidirectional communication between adaptive and innate immune cells within vulnerable brain regions after BmΔvjbR strain treatment driven by expression of tnaA. Whether this reflects enhanced antigen presentation by microglia, altered T-cell effector function, or both remain unclear.

It is also possible that we observed a floor effect of exacerbated neurodegeneration in this model by BmΔvjbR::tnaA treatment such that the ∼20% increase in tissue atrophy observed compared to the PBS-treated TE4 group was insufficient to observe significant changes in cytotoxic CD8^+^ T cell populations that have been more directly implicated as a cause for neurodegeneration in this model (6) and in the cerebrospinal fluid of individuals with Alzheimer’s Disease (4). Future studies should aim to address these possible mechanisms via characterization of peripheral and brain immune cell population changes via flow cytometry or single-cell RNA sequencing on CD45 high-enriched populations (5). In sum, these findings support a model in which the neurodegenerative consequences of peripheral immune modulation are mediated through qualitative changes in neuroimmune interactions rather than broad changes in initial events of innate immune activation within the brain that respond to tau pathology.

A key limitation of this study is that we have not characterized peripheral and infiltrating T cell subsets, particularly CD4^+^ T cells that are pro-inflammatory (Th1, Th17) or anti-inflammatory (regulatory T cells). Th17 responses have been shown to influence cognitive decline induced by dietary salt hypertension (26,27), but our peripheral cytokine analysis suggests little changes in circulating levels of IL-17. The BmΔvjbR::tnaA strain was originally developed to support the production and function of regulatory T cells, and the strongest effects of BmΔvjbR::tnaA treatment were observed in combination with adoptive regulatory T cell transfer in a collagen-induced arthritis model of autoimmunity (16). Although regulatory T cells that reside in the dural meninges support brain homeostasis (28), depletion of regulatory T cells in a model of Aβ amyloidosis improved pathological phenotypes (29). These observations support the idea that disease context is critical for manipulation of these cell types for therapeutic benefit.

These data also highlight the possible challenges of using immune modulation with bacterial therapeutics at a stage of disease where significant pathology has developed and neurodegenerative processes have been initiated. In our study, we selected a time point to begin intervention, 6 months, near the onset of tau pathology. In humans, Aβ accumulation begins in individuals decades before the onset of cognitive impairment but the onset of cortical tau pathology in individuals with AD begins only a few years before cognitive symptoms appear (30). Interestingly, results suggesting reduced dementia incidence in individuals vaccinated for herpes zoster or varicella zoster viruses (10–13) at advanced age (70-79 years or before the age of 80) raise the possibility that modifying the immune system in prodromal stages of dementia late in life is a viable therapeutic strategy. Future studies could adopt different administration strategies, such as prevention of pathology with a single administration of these bacterial strains earlier in life to reflect a scenario of identifying patients that are negative for biomarkers of tau pathology to achieve maximum benefits of preventing neurodegeneration and cognitive impairment(31).

## Conclusion

Collectively, these findings demonstrate that an engineered bacterial therapeutic with reported immunoregulatory activity may be disease context dependent. More broadly, the results suggest that the consequences of peripheral immune modulation via attenuated living pathogens during tauopathy may depend on how interventions reshape interactions between adaptive and innate immune cells within the CNS.

## List of Abbreviations

AD: Alzheimer Disease
Aβ: amyloid-beta peptide
Tau: microtubule-associated protein tau
CNS: Central Nervous System
*BmΔvjbR*: *Brucella melitensis* 16M harboring a deletion in vjbR, a master regulator of virulence
tnaA: Tryptophanase A
TE4: PS19 tau transgenic mice overexpressing human P301S tau by a mouse prion protein promoter on a human APOE4 knock-in background
IBA1: Ionized Calcium-binding Adapter Molecule 1
MHC-II: Major Histocompatibility Complex Class II
P2RY12: Purinergic receptor P2Y12
Th1: Type 1 Helper T cells
Th17: T helper 17 cells

## Declarations

### Availability of data and materials

All data are available in the main text or the additional materials.

### Competing Interests

WKS consults for Astonishing Labs.

PDF has equity in Tranquility Biodesign, ROXO Biotechnology, HelioWave Technologies, Suho Biotechnology, and SUBI Biotechnology.

DMH has equity in and is on the scientific advisory board of C2N Diagnostics; is on the scientific advisory boards of Denali, Genentech, Acta, and Switch Therapeutics; and consults for Pfizer, Novartis, Annexon, and Roche.

GG, PS, NP, YZ, MEB, CA, JDU, and JS have no interests to disclose.

### Funding

This research was funded by the Cure Alzheimer’s Fund (DMH) and the Research Education Component of NIH Grant Number 5P30AG06644407 (WKS). It was also funded, in part, by ARPA-H (No. 1AY1AX000010-01), NIH (R01CA273002) and the Roy Blunt NextGen Precision Health Endowment at the University of Missouri to PDF. The content of the presented information does not necessarily reflect the position or the policy of the Government, and no official endorsement should be inferred.

### Authors’ Contributions

DMH, PDF, CA and WKS conceived and designed experiments. WKS, GG, PS, AA, MB, YZ, and NP performed experiments and acquired data. WKS, YZ, CA, JDU, PDF, JS and DMH interpreted the data. WKS wrote the first draft of the manuscript, with revisions given by all authors.

## Acknowledgements

We thank Austin Allison for technical assistance with this study. We thank the Hope Center Alafi Neuroimaging Lab at Washington University School of Medicine for images acquired on the Nanozoomer scanner (NIH Shared Instrument Grant S10 RR027552). We thank Dr. Young Ah Goo, Minsoo Son, Jaenyeon Kim, and the Mass Spectrometry Technology Access Center at the McDonnell Genome Institute (MTAC@MGI) at Washington University School of Medicine, supported by the Diabetes Research Center/NIH grant P30 DK020579, Institute of Clinical and Translational Sciences/NCATS CTSA award UL1 TR002345, and Siteman Cancer Center/NCI CCSG grant P30 CA091842 for assistance with untargeted metabolomics analysis. We thank Dr. Diane Bender for assistance with plasma cytokine immunoassays. This work was supported in part by the Bursky Center for Human Immunology and Immunotherapy at Washington University, Immunomonitoring Laboratory.

## Additional Figures

**Additional Figure 1:**
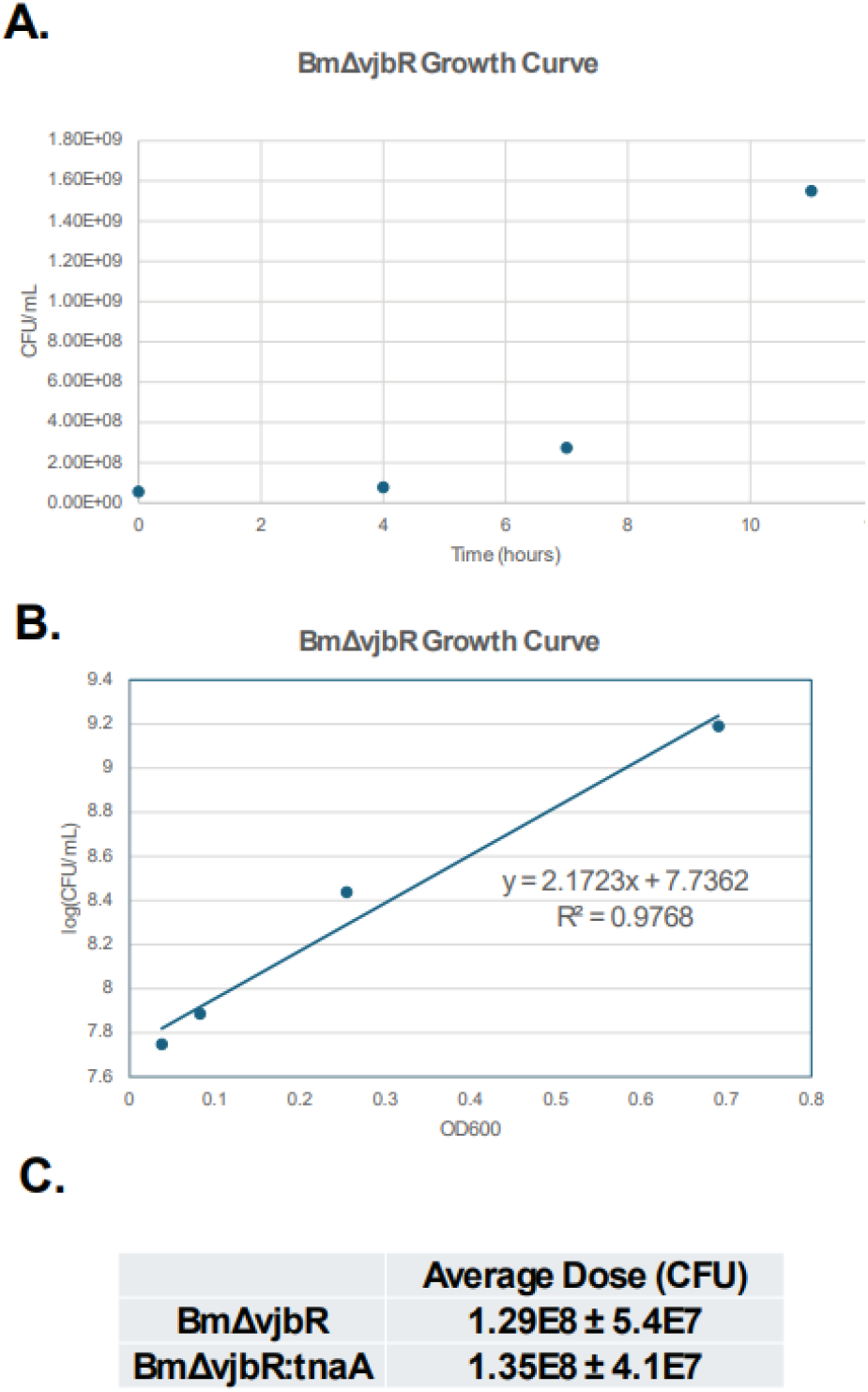
BmΔvjbR growth curves used for calculating treatment dosages. A.), B.) Growth curves used to calculate CFU/mL or each BmΔvjbR strain culture preparation. C.) Calculated average dose across all bacterial culture preparations for intraperitoneal injections for each strain.

**Additional Figure 2.**
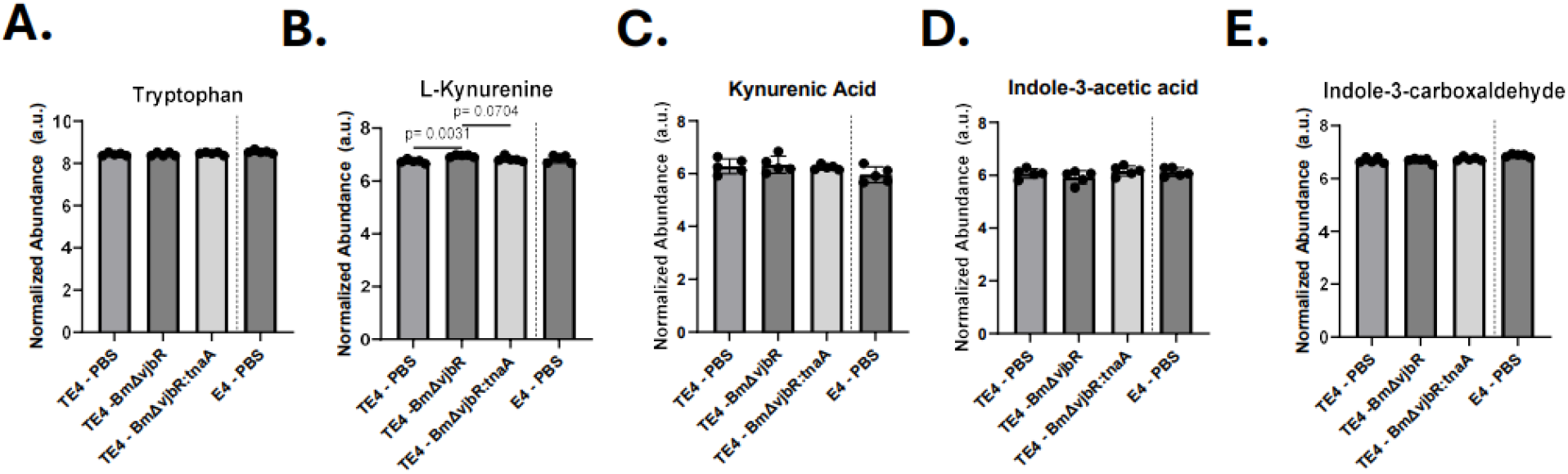
No differences between Brucella-treated groups in abundance of select tryptophan catabolites in plasma (Related to Figure 1). Quantile-normalized abundances of plasma indoxyl sulfate levels in end-stage, 285-day old mice. 1-way ANOVA on TE4-treated animals: F: 14.71 (2,12), p: 0.006. N = 5 animals per TE4-treated group for metabolomic analysis. P-values for individual comparisons with Tukey’s multiple comparisons test *post hoc*. Bar height represents mean value; error bars represent standard deviation.

**Additional Figure 3.**
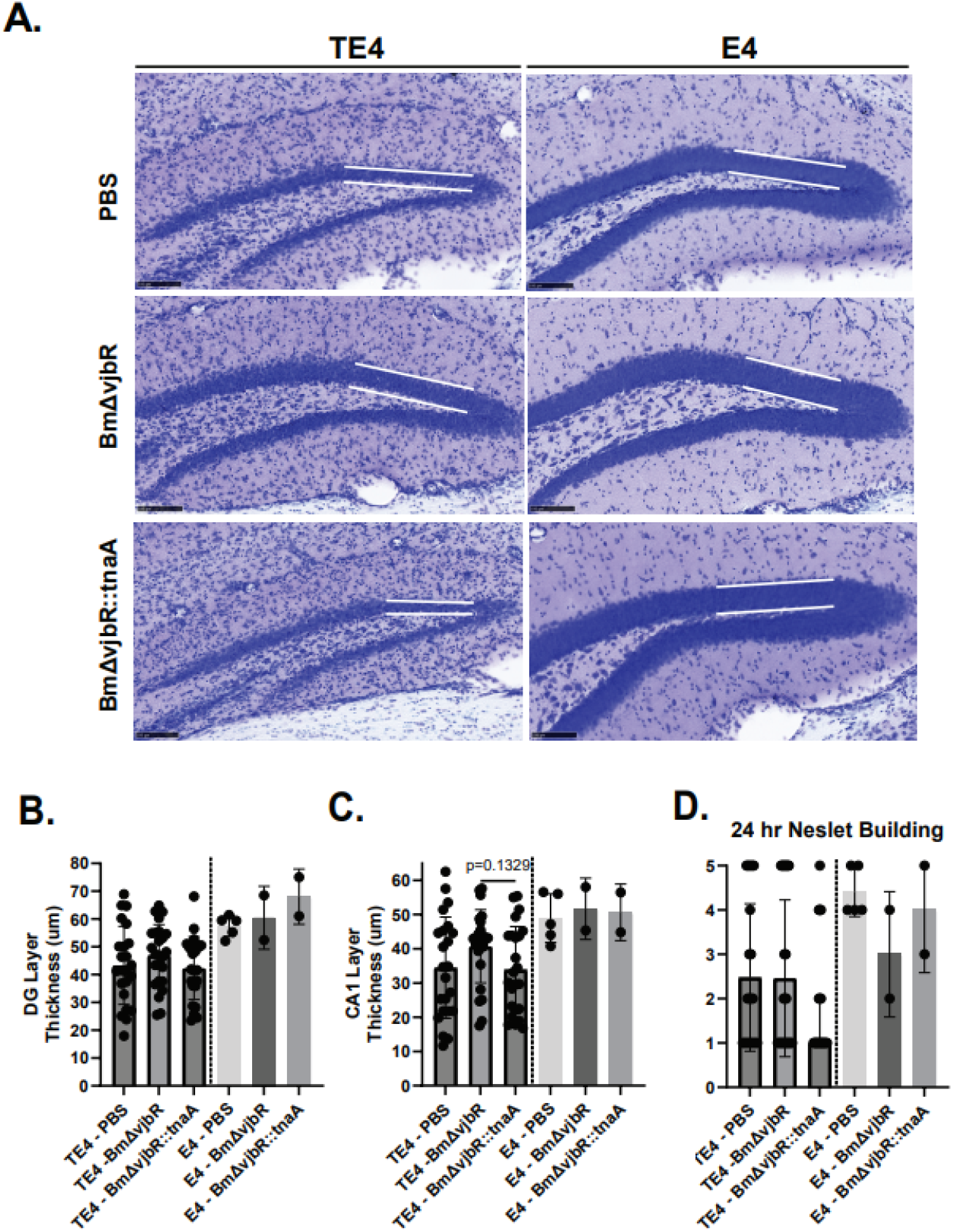
Repeated BmΔvjbR::tnaA treatment exacerbates tau-mediated neurodegeneration (related to Figure 2). A.) Representative image of granular cell layer thickness in the dentate gyrus. Scale Bar: 100 um. B.) Quantification of dentate gyrus (DG) cell layer thickness. F: 0.9689 (2,69), p-value = 0.3846. C.) CA1 pyrimadal cell layer thickness. W: 2.541 (2.00, 44.52), p-value: 0.0902. D.) Quantification of 24 hour nestlet building behavior prior to euthanasisa. Kruskal-Wallis test p-value: 0.1715. Number of animals analyzed in TE4 groups: PBS: 23, BmΔvjbR: 25, BmΔvjbR::tnaA: 24. F statistic denotes 1-Way ANOVA test and W statistic denotes Welch’s ANOVA test for TE4-treated groups. P-values for individual comparisons on graphs represent results of Tukey’s multiple comparisons test or Dunnett’s T3 multiple comparisons test *post hoc*. For B.) and C.) Bar height represents mean value; error bars represent standard deviation. For D.) Bar height represents median and error bars represent 95% confidence interval. Number of animals analyzed in TE4 groups: PBS: 23, BmΔvjbR: 25, BmΔvjbR::tnaA: 24.

**Additional Figure 4.**
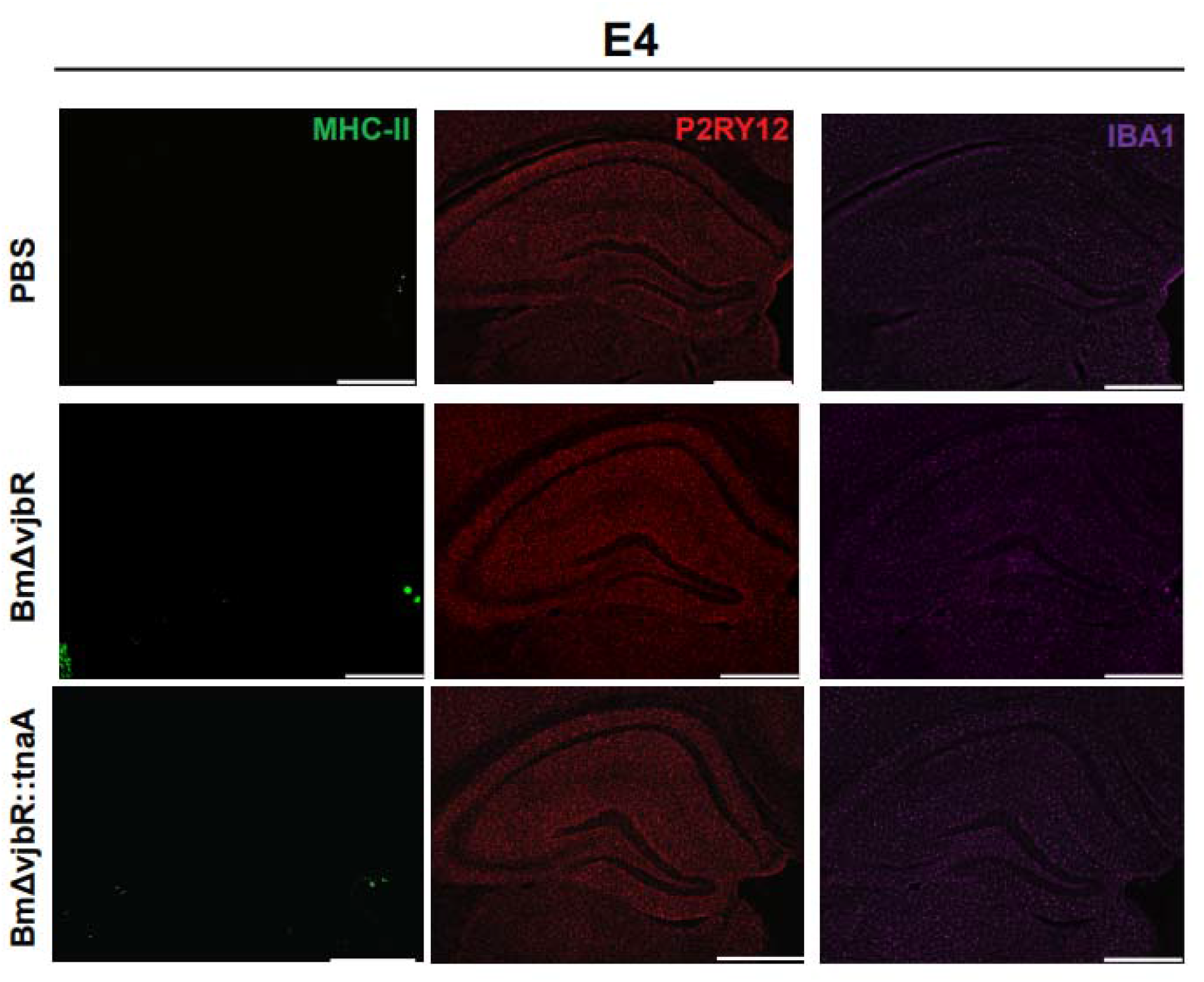
Repeated BmΔvjbR::tnaA treatment increases degeneration-associated microglial reactivity in TE4 mice (related to Figure 4). Representative images of hippocampal reactive microgliosis markers in 285-day-old nontransgenic E4 mice. Scale bar: 500 um.

**Additional Figure 5.**
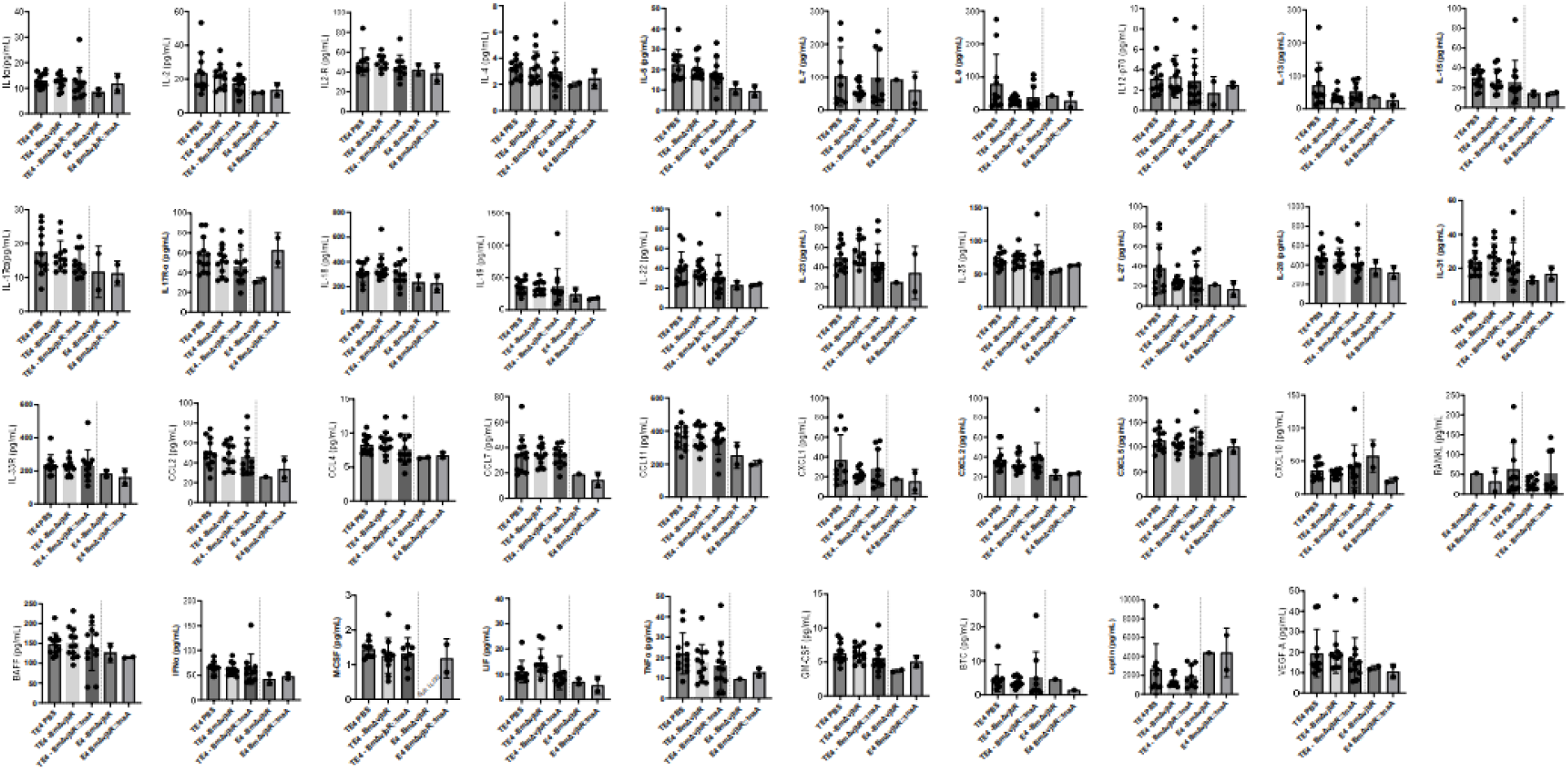
Remaining plasma cytokines, chemokines, and growth factors assayed, related to Figure 6. Abundances of all analytes measured that did not reach statistical significance. Statistics performed were either 1-way ANOVA or Welch’s ANOVA on TE4-treated groups. Bar height represents mean value; error bars represent standard deviation. For M-CSF measurements, samples in the E4-BmΔvjbR group were not detected above the lower limit of quantification, denoted by “Sub-LLOQ”. N = 10-12 per TE4-treated group.

